# A novel feature fusion based on the evolutionary features for protein fold recognition using support vector machines

**DOI:** 10.1101/845727

**Authors:** Mohammad Saleh Refahi, A. Mir, Jalal A. Nasiri

**Affiliations:** Department of Electrical Engineering, Amirkabir University of Technology, Tehran, Iran; Iranian Research Institute for Information Science and Technology (IranDoc), Tehran, Iran

**Keywords:** Protein Fold Recognition, Feature fusion, Evolutionary method, IG, Support Vector Machine

## Abstract

Protein fold recognition plays a crucial role in discovering three-dimensional structure of proteins and protein functions. Several approaches have been employed for the prediction of protein folds. Some of these approaches are based on extracting features from protein sequences and using a strong classifier. Feature extraction techniques generally utilize syntactical-based information, evolutionary-based information and physiochemical-based information to extract features. In recent years, Finding an efficient technique for integrating discriminate features have been received advancing attention. In this study, we integrate Auto-Cross-Covariance (ACC) and Separated dimer (SD) evolutionary feature extraction methods. The results features are scored by Information gain (IG) to define and select several discriminated features. According to three benchmark datasets, DD, RDD and EDD, the results of the support vector machine (SVM) show more than 6% improvement in accuracy on these benchmark datasets.

## 1. Introduction

Proteins are Jack of all trades biological macromolecules. They are involved in almost every biological reaction; Protein plays a critical roll in many different areas such as building muscle, hormone production, enzyme, immune function, and energy.

Typically more than 20,000 proteins exist in human cells, to acquire knowledge about the protein function and interactions, the prediction of protein structural classes is extremely useful [1]. Fold recognition is one of the fundamental methods in protein structure and function prediction.

Protein can be demonstrated as a chain of amino acids. Proteins with unique lengths and similarities are part of the same fold. They also have identical protein secondary structure in the same topology. Certainly, they have a regular origin [2].

One of the main steps which can be assumed as a vital stage for predicting protein fold is feature extraction. Computational feature extraction methods are divided into syntactical, physiochemical and evolutionary methods. Syntactical methods pay attention only to the protein sequence, like composition and occurrence [3, 4]. Physiochemical methods consider some physical and chemical properties of protein sequences. Evolutionary methods extract features from Basic Local Alignment Search Tool(BLAST).

When attempting to solve many biological problems it is obvious that a single data source might not be informative, and combining several complementary biological data sources will lead to a more accurate result. When we study methods of protein fold recognition, we found that less attention has been paid to the fusion of features to get more comprehensive features. In recent studies, researchers attempted to find new feature extraction methods[[5, 6, 7, 8, 9]] or train different classifiers to achieve high accuracy[[10, 11, 12, 13]], even though some problems like incomplete data sources, false positive information, multiple aspect problem,… encourage us to combine data sources.

Hence, to prepare more informative and discriminative features, we use Auto-Cross-Covariance(ACC)[8] and Separated dimer(SD)[7] methods. Because SD explores some amino acid dimers that may be non-adjacent in sequence [7] and ACC method measures the correlation between the same and different properties of amino acids [8]. One of the main advantages of ACC and SD is to find a fixed length vector from a variable protein length. The performance of the proposed method is evaluated using three benchmark datasets DD[3], RDD[14] and EDD[8].

In this paper, we focus on fusing ACC and SD feature extraction methods based on Position Specific Scoring Matrix(PSSM) generated by using the Position-Specific Iterated BLAST(PSI-BLAST) profile to predict protein fold.

The 1600 ACC features and the 400 SD features are extracted based on the PSSM. Finally, we construct a reduced-dimensional feature vector for the Support Vector Machine (SVM) classifier by using the Information Gain(IG).

The remaining sections of the paper are organized as follows. Section 2 describes the related works of the existing techniques. The methodology is explained in Section 3. Section 4 shows the experimental results and discussion. Finally, the conclusion and future works are given in Section5.

## 2. Related work

In 1997, Dubchak et al. studied syntactical and physiochemical method [15]. In which they assumed five properties of amino acid like hydrophobicity (H), frequency of *α* helix (X), polarity (P), polarizability (Z) and van der Waals volume (V). In [16] Forward Consecutive Search (FCS) scheme which trains physiochemical attributes for protein fold recognition.

Another solution to find similarity between protein sequences is based on the BLAST. Many feature extraction methods use BLAST alignment to extract the possibility of amino acid in specific positions called as PSSM. In 2009, pairwise frequencies of amino acids separated by one residue (PF1) and pairwise frequencies of adjacent amino acid residues (PF2) were proposed by Ghatny and Pal [5]. The bigram feature extraction method was introduced by Sharma et al. [6], bigram feature vector is computed by counting the bigram frequencies of occurrence from PSSM. In 2011, combination of PSSM with Auto Covariance (AC) transformation, was introduced as feature extraction method [17].

Another method introduced by Saini et al. [7] is separated dimers(SD); they used probabilistic expressions of amino acid dimer occurrence that have varying degrees of spatial separation in the protein sequence to predict protein fold. Dong et al. [8] proposed autocross-covariance (ACC) transformation for protein fold recognition. Moreover, Pailwal et al. [9] proposed the ability of trigram to extract features from the neighborhood information of amino acid.

In addition to the feature extraction methods, some researchers have paid attention to classification methods for protein fold recognition. In [10] Kohonens selforganization neural network is used and showed the structural class of protein is considerably correlated with its amino acid composition features.

Baldi et al.[18] employed Recurrent and Recursive Artificial Neural Networks (RNNs) and mixed it by directed acyclic graphs (DAGs) to predict protein structure.

In [12], classwise optimized feature sets are used. SVM classifiers are coupled with probability estimates to make the final prediction. Linear discriminant analysis(LDA) was employed to evaluate the contribution of sequence parameters in determining the protein structural class. Parameters were used as inputs of the artificial neural networks[19]. The composition entropy was proposed to represent apoptosis protein sequences, and an ensemble classifier FKNN (fuzzy K-nearest neighbor) was used as a predictor[13].

TAXFOLD [20]method extract sequence evolution features from PSI-BLAST profiles and the secondary structure features from PSIPRED profiles, and finally a set of 137 features is constructed to predict protein folds.

Sequence-Based Prediction of ProteinPeptide(SPRINT) method is used to the prediction of Proteinpeptide Residue-level Interactions by SVM [11]. SVM implements the structural risk minimization (SRM) that minimizes the upper bound of generation error [21, 22].

The DeepCov method proposed in [23], this method uses convolutional neural networks to work on amino acid pair frequency or covariance data extract from sequence alignments.

In [24] is attempted to show Artificial Neural Network (ANN) with different feature extraction method is more accurate than other classifier methods.

## 3. Methodology

This section illustrates the step-by-step of the proposed method for protein fold recognition. In the first step, sequence alignments are found for each protein using BLAST. To show improvements in protein fold recognition using evolutionary information that are presented in PSSM, therefore ACC [8]and SD [7] features are extracted from PSSM. In the next step, the features are combined and selected by the IG. In the last step, the SVM algorithm is trained to classify proteins. A comprehensive view of this approach can be found in Figure1.

**Figure 1:**
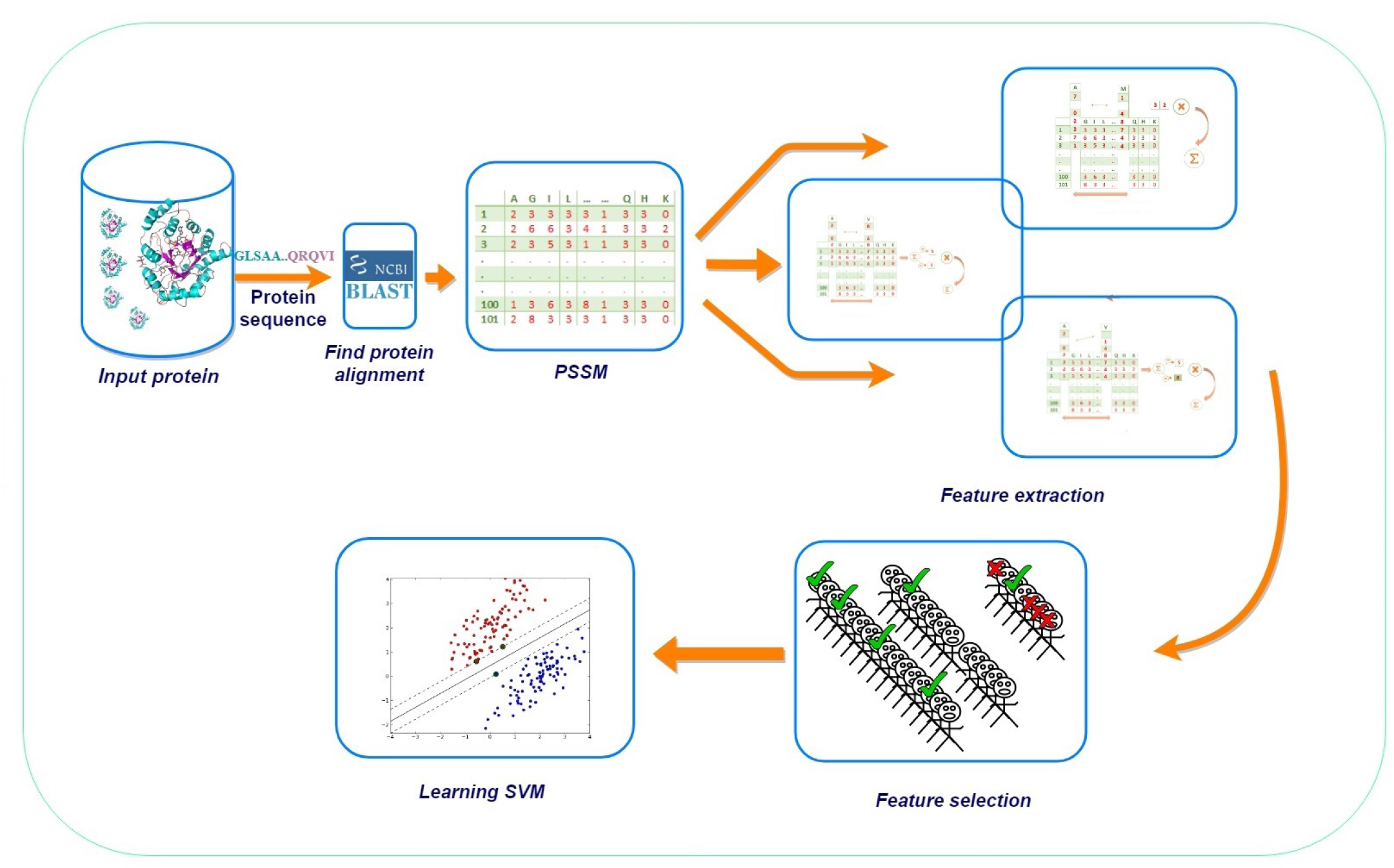
The flowchart of the proposed method pipeline. Sequence alignments are found for each protein by BLAST. PSSM is calculated to extract the ACC and the SD.The features are selected by the IG. The SVM algorithm is trained to classify proteins.

### 3.1. Preprocessing

#### 3.1.1. BLAST

Similarity is used here to mention the resemblance or percentage of identity between two protein sequences [25]. The similarity search depends on the bioinformatics algorithm. Basic Local Alignment Search Tool(BLAST) is a tool that helps researchers to compare a query sequence with a database of sequences and identify specific sequences that resemble the query sequence above a certain threshold. BLAST is a local alignment algorithm that means to find the region (or regions) of the highest similarity between two sequences and build the alignment outward from there [26].

#### 3.1.2. PSSM

Position Specific Scoring Matrix(PSSM) is used to express motif in a protein sequence. P-BLAST searches in which amino acid substitution scores are given separately for each position in a protein multiple sequence alignment. In this paper, PSSM is used to extract features by ACC and SD methods.

### 3.2. Feature Extraction

#### 3.2.1. ACC

ACC fold [8] utilizes autocross-covariance transformation that convert the PSSMs of different lengths into fixed-length vectors. The ACC separates two kinds of features: AC between the same properties, cross-covariance (CC) between two different properties. The AC variable measures the correlation of the same property between two properties separated by LG, distance along the sequence:

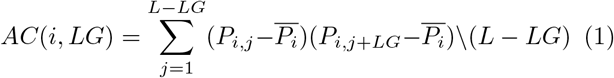

Where *P*_*i,j*_ is the PSSM score of amino acid i at position j, and 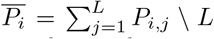, the average score of an amino acid i in the total protein sequence. The number of features which are calculated from AC is 20 * *LG*. The CC measures the correlation of two different properties between the distances of LG along the sequence:

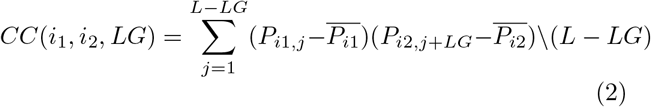

The CC variables are not symmetric. The total number of CC variables is 380 * *LG*.The combination of AC and CC features make 400 * *LG* feature vectors.

#### 3.2.2. SD

Separated Dimer(SD) method was introduced by Saini et al.[7]. It attempts to extract features from amino acids that may or may not be adjacent in the protein sequence. The SD demonstrates the probabilities of the occurrence of amino acid. SD generates 400 features.

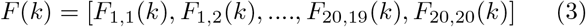

*F* (*k*) is computed as the feature sets for probabilistic occurrence of amino acid dimers with different values of *k* which is a special distance between dimers. It is obvious if *P* is in the PSSM matrix for a protein sequences, it is *L*×20 matrix where L is the length of the protein sequence:

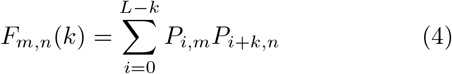

in which *m, n* (1 ≤ *m, n* ≤ 20) is the score of two selective amino acid in PSSM.

### 3.3. Fusion hypothesis

More attention needs to be paid to find an efficient technique for integrate distinct data sources for the protein fold recognition problem[27]. Various techniques have been employed based on the features which are extracted from protein sequences.

These techniques investigate different aspects of a sequence like the study of possible position of amino acids, protein chemical characteristics and syntactical features …. Hence, integrating them can model the folding problem more accurate.

In this study three hypotheses have been considered for fusion data sources. The first, only evolutionary features are used since integrating different types of features may have an undesirable effect on each other.

The next assumption is considered choosing the ACC and SD methods. When we studied the recent paper, we observed the recall and precision of some protein folds which were almost equal to high value or one was a complement of the other. Hence, when the recall of the ACC(SD) is low, then the recall of the SD(ACC) is high, and also for the precision, we observe this behavior, in almost every fold.

The last hypothesis is that the ACC and the SD features showed a relationship between amino acids which may or may not be adjacent. In this approach, three different characters are defined which show each amino acid in a specific position what relation has with others. These characters are shown in Figures23,4.

**Figure 2:**
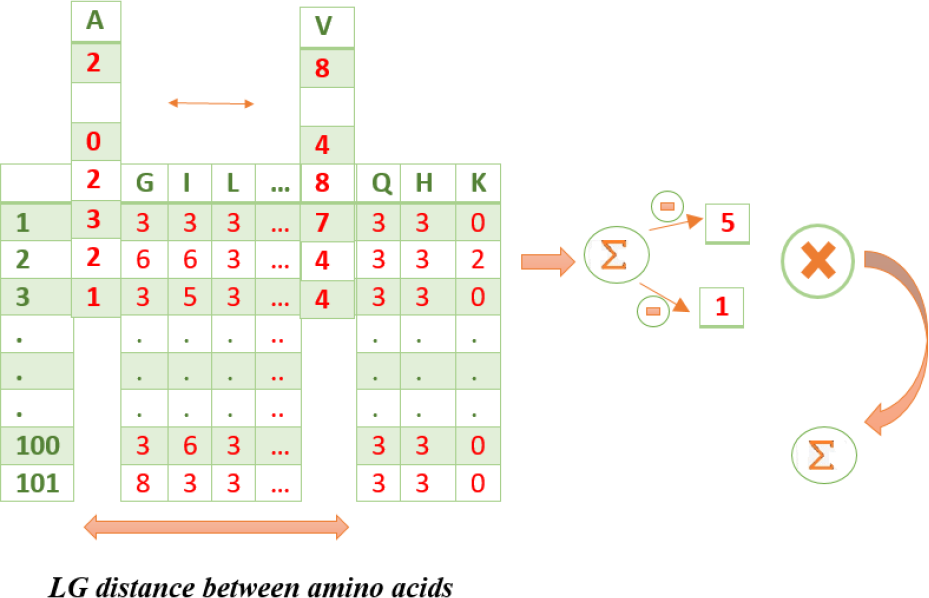
The AC features of the ACC, measures the correlation of the same property between two properties separated by a distance of LG along the sequence

**Figure 3:**
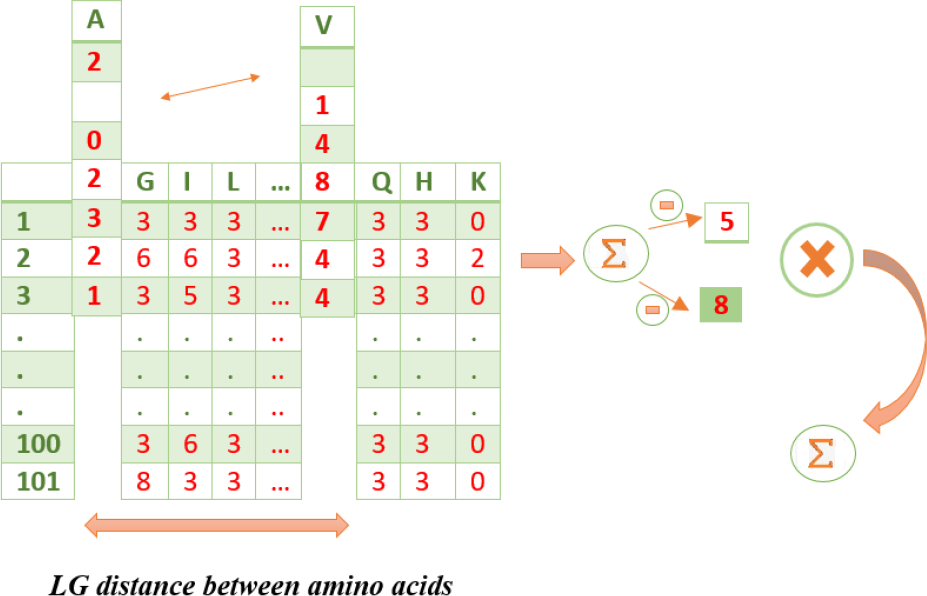
The CC features of the ACC, measures the correlation of two different properties between the distances of LG along the sequence

**Figure 4:**
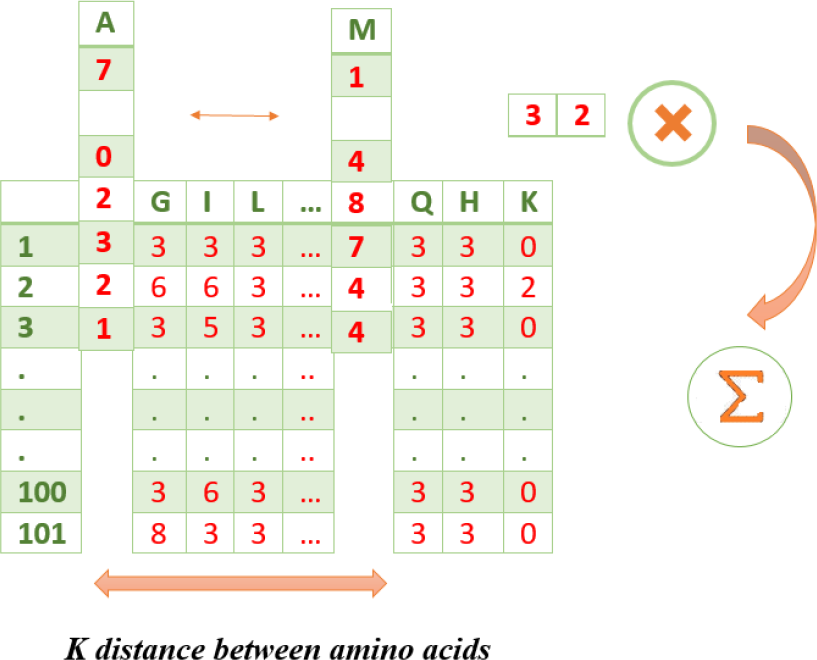
The SD consist of aminoacid dimers with probabilistic expressions that have k separation.

### 3.4. Information Gain

Feature selection is a common stage in classification problems. It can improve the prediction accuracy of classifiers by identifying relevant features. Moreover, feature selection often reduces the training time of a classifier by reducing the number of features which are going to be analyzed.

Information gain (IG) is a popular feature selection method. It ranks features by considering their presence and absence in each class [28]. The IG method gives a high score to the features that occur frequently in a class and rarely in other classes.

Given *T* the set of training samples, *x*_*i*_ the vector of *i*th variables in this set, 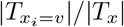 the fraction of samples of the *i*th variable having value *v*. The IG method can be computed as follows:

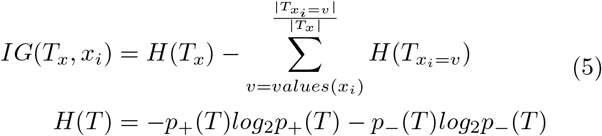

where *p*_±_ denotes the probability of a sample in the set *T* to be of the positive or negative class.

### 3.5. Support Vector Machine

Support Vector Machine (SVM) was proposed by Vapnik and Cortes in 1995 [29]. It is a powerful tool for binary classification. SVM is on the basis of Structural Risk Minimization (SRM) and Vapnik-Chervonenkis (VC) dimension. The central idea of SVM is to find the optimal separating hyperplane with the largest margin between the classes. Due to the SRM principle, SVM has great generalization ability. Moreover, the parameters of the optimal separating hyperplane can be obtained by solving a convex quadratic programming problem (QPP), which is defined as follows:

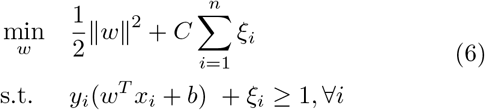

where *ξ* is the slack variable associated with *x*_*i*_ sample and *C* is a penalty parameter. Note that the optimization problem can be solved when the classification task is linearly separable. In the case of nonlinear problems, the input data is transformed into a higher-dimensional
feature space in order to make data linearly separable. It makes possible to find a nonlinear decision boundary without computing the parameters of the optimal hyperplane in a high dimensional feature space [30].

As mentioned in this subsection, SVM is designed to solve binary classification problems. However, there are multi-class approaches such as One-vs-One (OVO) and One-vs-All (OVA) [31], which can be used for solving multi-class classification problems. In this paper, we used OVO strategy.

## 4. Experimental Result

### 4.1. Dataset

Three popular datasets which were employed in this study are DD dataset [3], EDD dataset [8], and RDD dataset[14]. DD dataset contains 27 folds which represent four major structure classes: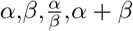. The training set and the testing set contains 311 training sequences and 383 testing sequences whose sequence similarity is less than 35%[3]. The EDD dataset consists of 3418 proteins with less than 40% sequential similarity belonging to the 27 folds that originally are adopted from the DD dataset. The RDD dataset consists of 311 protein sequences in the training and 380 protein sequences in testing datasets with a similarity lower than 37% [14].

### 4.2. Result

The experiments were performed on the benchmark datasets to evaluate the performance of the classification due to our fusion method. we also adopted the 10-fold cross-validation in this study, which has done by many researchers to examine predictive potency.

In this study LibSVM [32] with RBF (Radial Basis Function) as the kernel functions has been used. The C parameter was optimized by search between {2^−14^, 2^−13^, …, 2^13^, 2^14^} and also Γ parameter of RBF was considered between {2^−14^, 2^−13^, …, 2^13^, 2^14^}. The SVM was originally designed for binary data classification.This study used one-versus-one method to approach a multi-class classifier.

The details of the feature extraction method are explained in methodology, but it is important to know how far is assumed between aminoacids, for each ACC and SD methods. In developing the algorithm to extract features from PSSM, *LG* and *k* parameters have been assumed like ACC and SD papers values[7, 8]. We considered both *LG* and *k* equals to 4.

The IG [28] makes our method safe from noisy features. In this approach, we considered the features ranked between [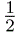 max_*IG*_, max_*IG*_] for each dataset. The results of IG for each dataset are exhibited in Table2.

### 4.3. Discussion

Table1 illustrates the total prediction accuracies of the existing approaches for classification of protein folds in the DD,RDD and EDD datasets. Table1 also shows the success rates of our proposed fusion approach. According to Table1, classification results of the combined ACC and SD followed by selection of best features by IG show considerable improvement compared to the state of art.

**Table 1:** Comparison of the proposed method with the existing predictor and Meta-predictors for the DD, RDD and EDD.

| Methods | Reference | DD | RDD | EDD |
| --- | --- | --- | --- | --- |
| occurrence | [4] | 42 | 56.6 | 70.0 |
| ACC | [8] | 68.0 | 73.8 | 85.9 |
| PF1 | [5] | 50.6 | 53.3 | 63.0 |
| PF2 | [5] | 48.2 | NA | 49.9 |
| TAXFOLD | [20] | 71.5 | 83.2 | NA |
| Bigram | [6] | 79.3 | 59.6 | 79.9 |
| SD | [7] | 86.3 | 72.1 | 90.0 |
| Trigram | [7] | 73.4 | 60.0 | 80.0 |
| MF-SRC | [33] | 78.6 | NA | 86.2 |
| Enhanced-SD | [24] | 90.0 | 75.4 | **93.0\*** |
| Proposed Method | - | **91.31** | **91.64** | 91.2 |

\* The evaluation method not defined in [24] approach.

**Table 2:** F1 score, Sensitivity and Precision, Measurement tools to evaluate the proposed method.

| Data set | F1 score | Sensitivity | Precision | Number of features |
| --- | --- | --- | --- | --- |
| DD | 0.98 | 0.92 | 0.93 | 1300 |
| RDD | 0.98 | 0.92 | 0.93 | 1416 |
| EDD | 0.96 | 0.91 | 0.93 | 900 |

Figure9 has been shown to figure out the result distribution of feature selection method. Even though the number of ACC in the three datasets is more but all of the SD features exist in selected features. However, we studied and compared SD and ACC methods separately, we found that the fusion of them can make more informative data which cover all characteristics of folds.

It is evident in Figure6, Figure7, and also Figure8, only “FAD-BINDING MOTIF” protein fold is not well recognized, and these confusion matrices show the power of proposed method for predicting the other folds in these datasets.

The Figure5 has been shown to evaluate the IG. It is obvious that maximum accuracy of classification for each dataset has been achieved when we consider ranking features higher than 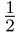 max_*IG*_ for these datasets.

**Figure 5:**
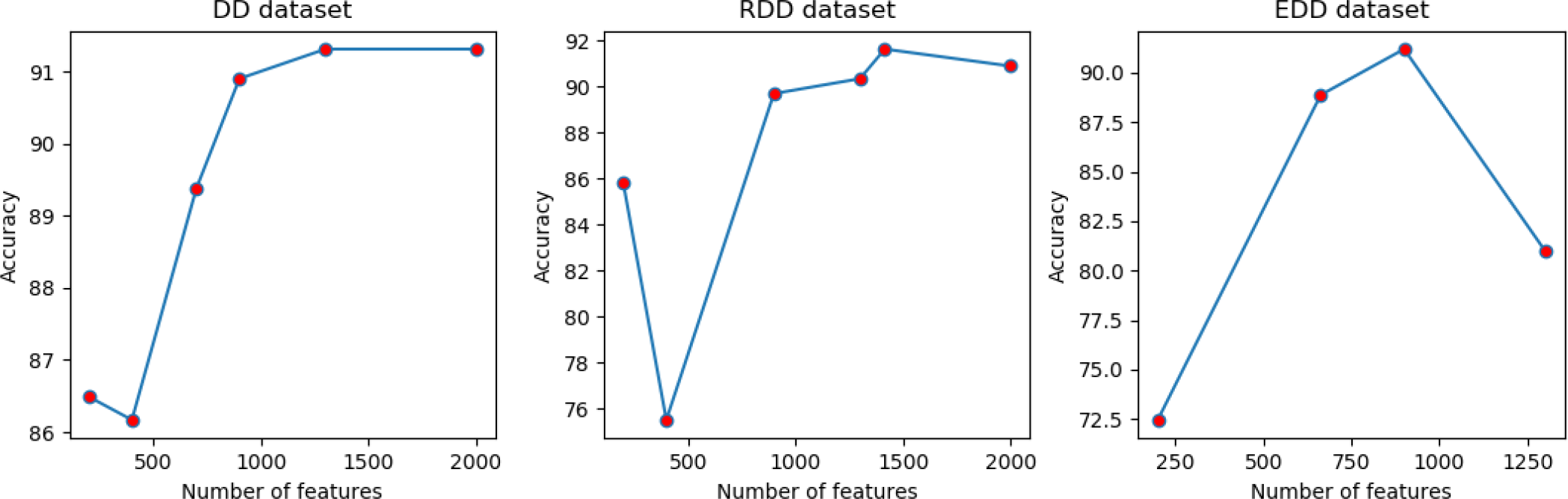
Comparison of number of features and accuracy for DD,RDD and EDD datasets to evaluate the IG method

**Figure 6:**
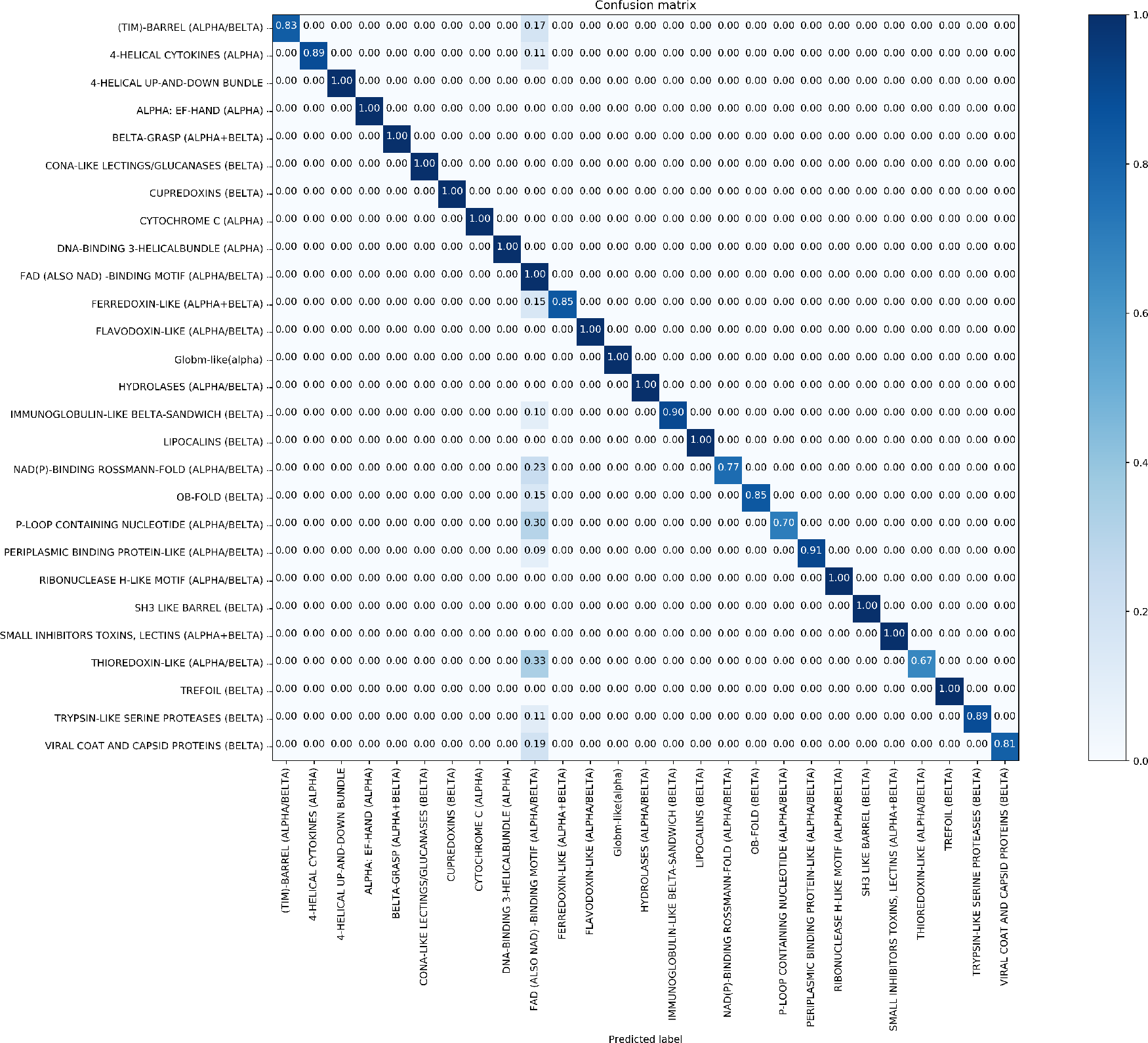
Confusion matrix of DD dataset(91.31%)

**Figure 7:**
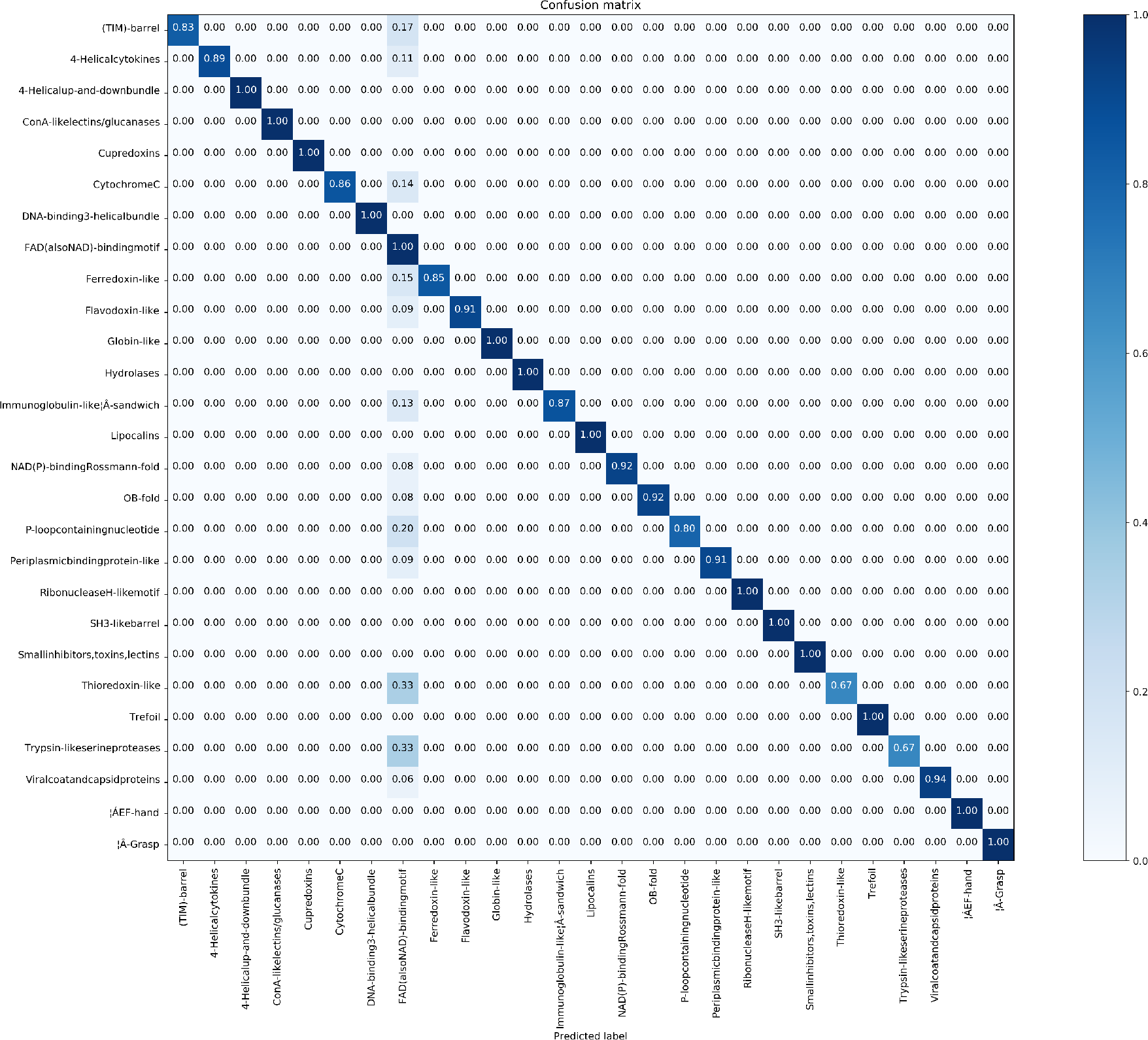
Confusion matrix of RDD dataset(91.64%)

**Figure 8:**
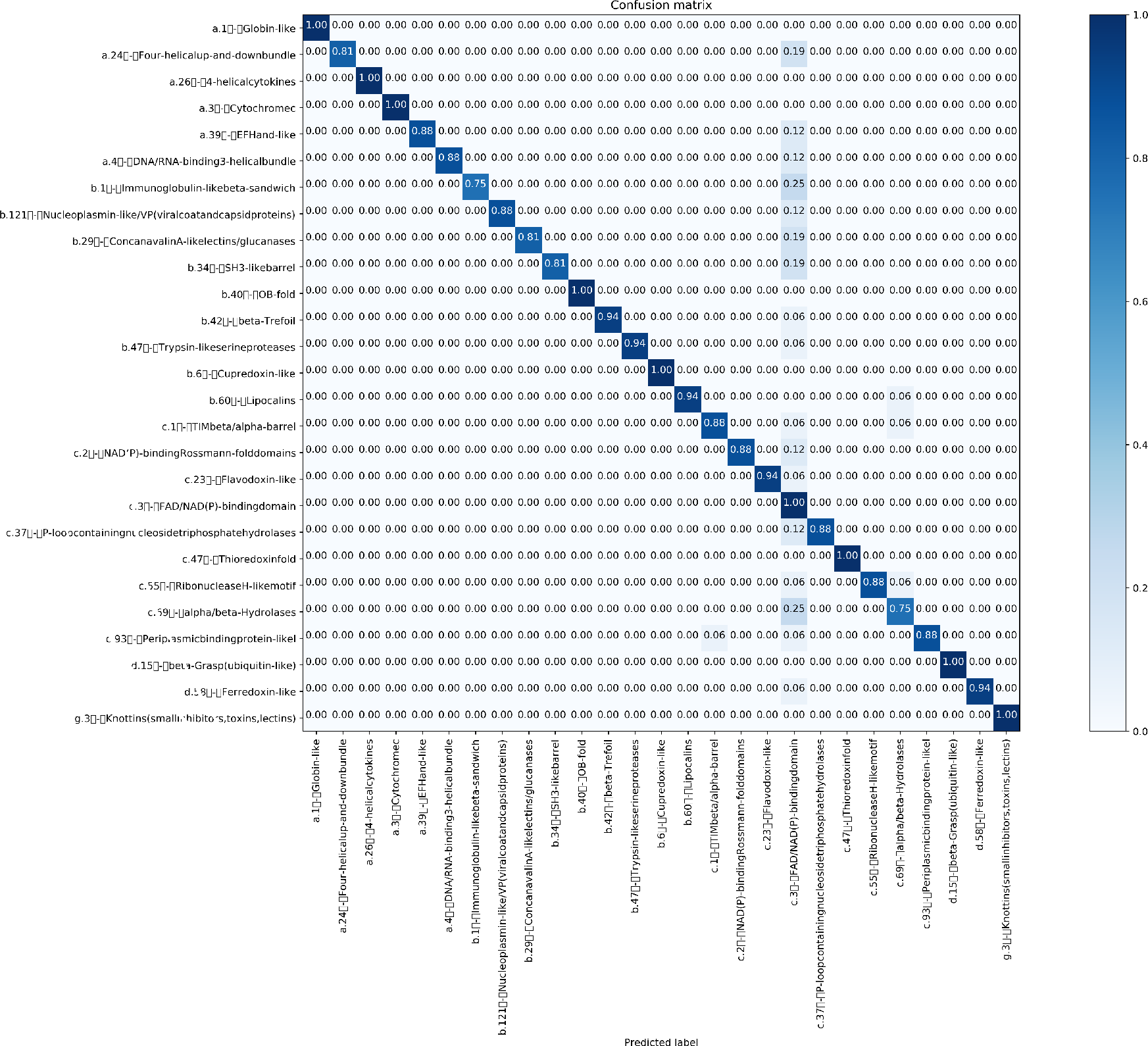
Confusion matrix of EDD dataset(91.2%)

**Figure 9:**
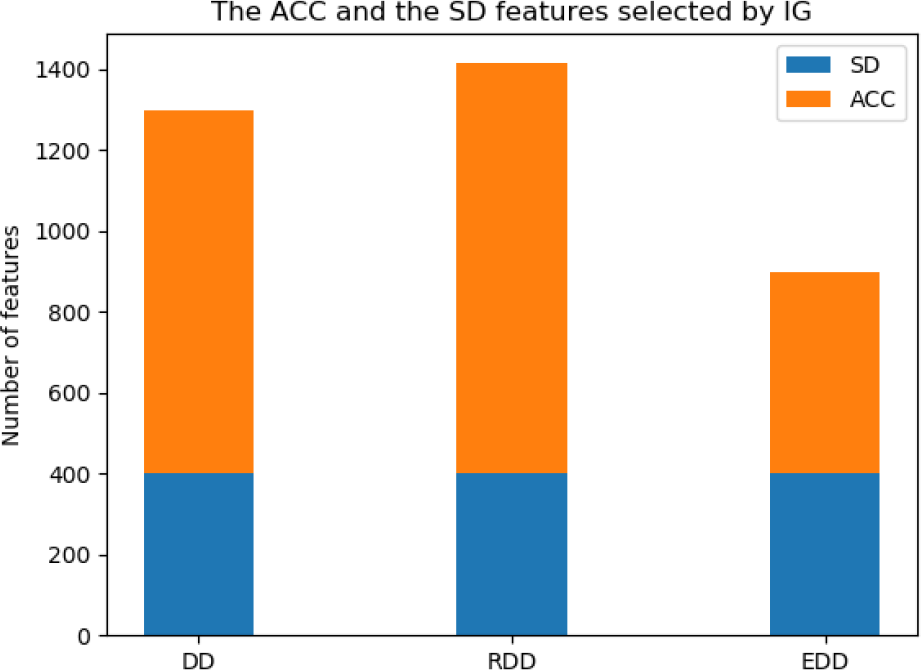
Comparison of the ACC and the SD in DD,RDD and EDD datasets.

Sensitivity measures the ratio of correctly classified samples to the whole number of test samples for each class which is classified as correct samples and calculated as follows:

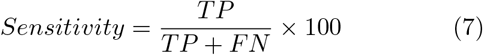

TP represents true positive and FN represents false negative samples. Precision represents, how relevant the number of TP is to the whole number of positive prediction and is calculated as follows:

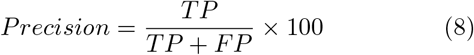

FP denotes false positive. F1 Score is the weighted average of Precision and Recall. F1 score, as other evaluation criteria which are used in this study measures, is calculated as follows:

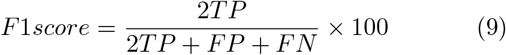

The sensitivity, precision, and F1 score are computed for each class and then averaged over all the classes which are calculated and published in Table2.

## 5. Conclusion

This study aims to improve protein fold recognition accuracy by fusing information that are extracted from the PSSM matrix. In this approach, we used ACC and SD feature extraction methods. It was observed that the proposed technique eventuates to 6% improvement for the accuracy of these three benchmark datasets.

In the future, classification can be done by combining more syntactical,physiochemical or evolutionary features.

To achieve more accuracy, future studies should be concentrate on “FAD-BINDING MOTIF” protein fold that has less discriminative features in the SD and the ACC. Boosting classifier may be employed to find better solutions for protein fold recognition.

## Notes

https://github.com/MsAlEhR/bio-protein-recog

